# Disordered glass nanowire substrates produce in vivo-like astrocyte morphology revealed by optical diffraction tomography

**DOI:** 10.1101/2024.11.14.623676

**Authors:** Pooja Anantha, Joo Ho Kim, Emanuela Saracino, Piyush Raj, Ivano Lucarini, Swati Tanwar, Jessica Chen, Luo Gu, Jay Agrawal, Annalisa Convertino, Ishan Barman

## Abstract

Astrocytes, integral components of the central nervous system (CNS), fulfill crucial roles such as maintaining ion homeostasis, providing neuroprotection, and contributing to the blood-brain barrier. Their distinctive, star-like morphology is essential to these functions, and abnormalities in astrocyte structure are linked to numerous neurological disorders. However, our understanding of astrocyte morphology, particularly *in vivo*, remains limited. Traditional imaging methods, such as fluorescence microscopy, introduce challenges like restricting continuous observation and comprehensive morphological analysis. In this study, we present a novel approach utilizing optical diffraction tomography (ODT), an advanced imaging technique that generates 3D refractive index profiles, to image and quantify detailed astrocyte morphology. We demonstrate, for the first time, the application of ODT to image samples through and on disordered glass nanowire (NW) substrates, overcoming the typical challenges posed by nanostructures, which can disrupt phase reconstruction. Crucially, we show that disordered glass nanowire (NW) substrates can induce *in vivo*-like astrocyte morphology in cultured rat cortical astrocytes. Compared to traditional glass substrates, astrocytes grown on disordered glass NWs substrates exhibited enhanced process branching and greater total arbor length—features typically observed in their natural, *in vivo* state, a state of advanced maturation. This finding underscores the significant influence of substrate topography on astrocyte structure and highlights the unique potential of nanostructured environments to mimic physiological conditions. By leveraging ODT, we were able to monitor astrocyte behavior on these substrates, providing unprecedented insights into their morphological dynamics. Our study pioneers the use of nanostructured substrates for reconstructing astrocyte morphology and sets the stage for further exploration of how microenvironmental cues shape astrocyte morphology and behavior.

## 1. Introduction

Understanding how living cells sense their environment and respond to it by adjusting their shape, migration, proliferation, advanced maturation potential and survival remains an open question relying on fundamental biophysical processes, including morphogenesis, cancer progression and tissue repair. Central to this understanding is the recognition that cell behavior within tissues is intricately linked to the mechanical properties of their surroundings, encompassing both neighboring cells and the extracellular matrix (ECM) [1], [2]. Mechanosensitive adhesion complexes at cell–substrate interfaces and cell–cell junctions orchestrate interactions, transducing physical signals critical for diverse cellular processes spanning multiple spatial-scales, from molecular to multicellular levels, and times-scales, from milliseconds to days [2], [3], [4], [5].

Nanostructured biomaterials offer a precise means of controlling and manipulating the mechanical environment at the nanoscale, making them powerful tools for studying mechanobiology. By simulating the intricate properties of the ECM, these nanomaterials provide essential platforms for investigating how cells adapt in response to controlled environmental cues [1], [6], [7], [8]. In this regard, nanomaterials can be used as supporting material for different biological purposes such as the study of cell adhesion, proliferation, structural and morphological features [9], [10], [11]. Particularly, nanostructures, such as nanopillars and nanowires, can influence cellular behavior by exerting mechanical forces that affect processes like adhesion, migration, proliferation, and advanced maturation. They have garnered significant interest in the field of neuroscience for their potential applications in studying neurodegenerative disorders (NDs) characterized by the progressive loss of specific brain population including Alzheimer’s disease (AD), Parkinson’s disease (PD), ischemia and epilepsy [12], [13], [14]. These nanomaterials allow for precise control over cell-substrate interactions, providing valuable insights into fundamental neurobiological processes and offering promising new avenues for diagnostic and therapeutic advancements. Despite their potential, many underlying mechanisms and phenomena associated with these interactions remain poorly understood.

Recent research underscores the potential of nanostructured materials that mimic the ECM to study cell-surface interactions at the nanoscale [7], [10]. These materials provide valuable insights into the mechanisms that govern neuronal development and function. Moreover, various studies have demonstrated how these nanostructures can act as scaffolds to guide neuronal growth and direct axonal outgrowth in a controlled manner, fostering specific neuronal responses and advancing our understanding of synaptic plasticity and neuronal circuitry in both healthy and diseased states [15], [16]. Additionally, nanowires can be arranged in arrays to serve as high-density electrodes, enabling the recording of electrical activity from individual neurons or neuronal networks with high spatial and temporal resolution. This capability holds promise for developing diagnostic tools to monitor disease progression or evaluate treatment efficacy [17], [18], [19], [20]. Over the past several decades, significant advancements in understanding brain networks have led to a growing focus on the critical role of non-excitable glial cells, particularly astrocytes. While astrocytes do not generate action potentials like neurons, they are vital for brain function, exhibiting bioelectrical activity facilitated by ion channel proteins that regulate the movement of ions and organic molecules across their membranes. This bioelectrical activity enables astrocytes to support and modulate neuronal activity, contributing to a wide range of processes essential for healthy brain function [21]. Disruptions in the structure and function of astrocytic ion channels contribute to neurodegenerative conditions and inflammation, leading to astrogliosis and central nervous system (CNS) dysfunction. [15], [22]. The expanding recognition of astrocytes’ roles in the CNS, including functions previously attributed solely to neurons—such as modulation of synaptic transmission and involvement in learning and memory—has fueled the demand for specialized tools to study these cells *in vitro*. A significant limitation of traditional 2D *in vitro* systems is that astrocytes lose their characteristic star-like morphology, which is crucial for their functional role in the brain. Instead, they adopt a flattened, polygonal shape, which does not accurately reflect their *in vivo* state or functionality [23]. This highlights the need for advanced *in vitro* models that can preserve the complex morphology of astrocytes, enabling more accurate studies of their behavior and interactions with other neural cells.

In parallel, the advancement of modern optical microscopy techniques has significantly improved our ability to visualize biological processes in their physiological context. Among these innovations, Optical Diffraction Tomography (ODT) stands out as a particularly powerful tool for studying cellular dynamics [24], [25], [26], [27], [28]. One of the key advantages of ODT is its ability to measure the three-dimensional refractive index (RI) distribution of biological samples, providing high-resolution 3D images of live cells. This advanced imaging technique enables the quantification of cellular characteristics with picogram precision, offering detailed insights into subtle changes in cell morphology, growth, and advanced maturation [24], [26], [28]. By capturing these cellular dynamics in 3D, ODT can reveal structural information that would be difficult to obtain with conventional 2D imaging techniques. When combined with nanostructured substrates, such as transparent glass nanowires (glass NWs), ODT becomes an even more powerful tool for studying cellular behavior. These substrates can mimic the mechanical properties of the ECM and provide nanoscale cues that guide cellular morphology and function. The transparency of the nanostructured substrates allows ODT to be used effectively, enabling the visualization of cells in a biomimetic environment that closely resembles their *in vivo* conditions. This combination allows for the precise study of how cells, including astrocytes, respond to nanoscale mechanical cues, while preserving their *in vivo*-like morphology and behavior. The ability to image living cells in real time on these substrates without perturbing the system offers unparalleled insights into cellular dynamics, providing a novel approach to investigate the complex roles of astrocytes in neurobiology and disease.

Here, we present, to the best of our knowledge, the first study combining transparent nanostructured substrates based on mats of transparent glass NWs with ODT to study the morphology of primary rat cortical astrocytes. This study is particularly significant given the increasing recognition of astrocytes’ pivotal role in brain function and pathology [29], [30], [31], [32]. In this study, we investigated the impact of glass NWs on the morphology of rat cortical astrocytes, aiming to understand how this substrate influences astrocyte behavior in comparison to traditional glass substrates. By culturing astrocytes on glass NWs substrates, we sought to create a biomimetic environment that encourages astrocytes to adopt *in vivo*-like morphological features. This approach allows for more accurate observations of astrocyte morphology, behavior, and interactions with the substrate, thereby providing a deeper understanding of their roles *in vivo*. The transparency of glass NWs substrates offers a unique advantage when combined with advanced imaging techniques like ODT. By utilizing a 2D nanowire array, we can observe the characteristic star-like morphology of astrocytes, including their delicate, branched processes radiating from the central cell body. This finding highlights the potential of nanostructured biomaterials as effective 2D platforms for investigating astrocyte behavior. **Figure 1** provides an overview of our study.

**Figure 1.**
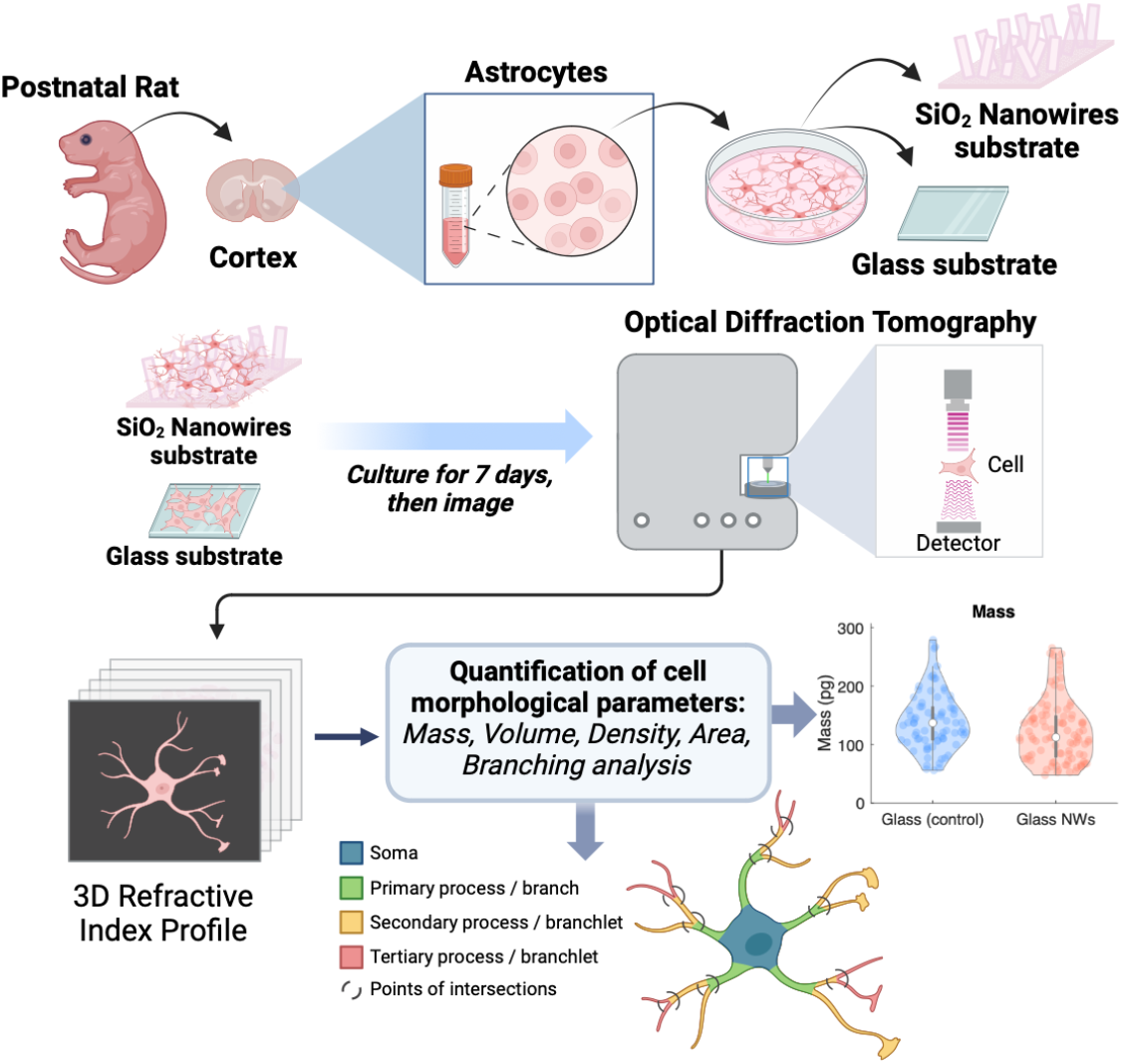
Schematic overview of the experimental workflow. Primary rat cortical astrocytes were cultured on transparent glass nanowire (NW) substrates and traditional glass substrates (control). The cells were imaged using Optical Diffraction Tomography (ODT) to assess their morphology. Morphological analysis was performed on the resulting 3D RI profiles of the astrocytes.

## 2. Methods

### 2.1. Fabrication of glass nanowires

SiO_2_ (glass) NWs were produced through thermal annealing of Si NWs grown by plasma enhanced chemical vapor deposition (PECVD). First, gold-induced Si NWs were grown on fused silica substrates [7], [33], [34], whose size fitted the holder stage of the ODT spectroscope. To induce the NWs growth, a 2 nm thick Au film was selectively evaporated onto the specific area, defined using photolithography and wet etching processes. The growth was performed with SiH_4_ and H_2_ as precursors at a total pressure of 1 Torr and substrate temperature of 350 °C. The flow ratio SiH_4_/(H_2_+SiH_4_) was fixed to 1:10. A 13.6 MHz radio frequency with power fixed at 5 W was used to ignite the plasma. The growth time was fixed at 7 min. A mat of dense and randomly oriented Si NWs, approximately long 2-3 mm, with an average diameter at the bottom of about 50– 80 nm, were obtained. After the growth, the Si NWs were thermally oxidized to form SiO_2_ (glass) NWs via a thermal treatment in a convection oven (controlled O2 atmosphere) at 980 °C for 8 hrs [35], [36], [37].

### 2.2. Scanning electron microscopy (SEM)

The morphology of the glass NWs mat was verified by a field emission scanning electron microscopy (FESEM) (ZEISS SIGMA 300) at an accelerating voltage of 5 kV. The length and diameter of the NWs were determined by using line tools of the image analysis program ImageJ. About 500 measurements of both diameter and length, obtained from 3 SEM images, were used to generate the related distribution for both as-grown Si NWs and glass NWs. The histograms were constructed using bin sizes that produced approximately 10 bins across the entire distribution.

### 2.3. Ultraviolet-visible (UV/Vis) spectroscopy

The optical properties were studied by measuring the transmittance and reflectance in the spectral range between 300 and 700 nm with a Lambda 35 UV–vis spectrophotometer.

### 2.4. Atomic force microscopy (AFM) characterization

AFM imaging of glass nanowire substrates was done on mica sheets of V1 quality using dry AFM. Bruker Multimode 8 scanning probe microscope with silicon cantilever (Bruker) was used for AFM imaging in tapping mode analysis.

### 2.4. Rat cortical astrocyte isolation and purification

All animal experimental procedures were conducted in strict accordance with the Animals Care and Use Committee at Johns Hopkins University. Astrocytes were extracted from postnatal day 2 Sprague-Dawley rats by initially isolating neural cell mixture from the rat followed by purification for astrocyte extraction [38], [39]. The meningeal layer was removed, and the cortex was dissected in cold HBSS (Thermofisher) with 2% P/S. The wet cortical tissue was then placed in a petri dish for chopping with a sterile blade. The chopped tissue (∼1mm^3^ pieces) was transferred to pre-warmed 0.05% trypsin-EDTA (Thermofisher) and incubated at 37°C shaken every 5 minutes. After 10 minutes, trypsin was quenched using 10% heat-inactivated fetal bovine serum (Cytiva), 100U/ml penicillin, and 100U/ml streptomycin (Thermofisher), and Dulbecco’s modified Eagle’s medium with glucose and pyruvate (DMEM, Thermofisher) solution (cDMEM). The tissue was triturated, centrifuged, and the pellet was resuspended in cDMEM. The cell suspension was further homogenized, strained through a cell strainer, and the process was repeated for a total of three times. Mixed neural cells were seeded onto poly-D-lysine (PDL, Thermofisher)-coated T75 culture flasks (one brain per flask). PDL coating involved incubating flasks with 10µg/mL PDL, diluted in deionized (DI) water, and added to each dish at a volume of 0.125mL/cm^2^ for 30 minutes at 37°C, followed by two washes with DI water for 5 minutes each. Initial seeding required 15 ml of culture medium, followed by 24 hours of culturing with subsequent 100% medium exchange to remove excess cell debris. The culture was maintained until cells reached over 90% confluence, which usually required a week, with media changes every 3 days. When cells were ready for purification, the flask was sealed, covered, and shaken on an orbital shaker at 210 rpm for 1 hour in a 37°C incubator then vigorously shaken around 10 times to dislodge remaining microglia and oligodendrocytes. The shaking focused on the flask’s bottom to ensure detachment. Following shaking, media was aspirated, monolayers were washed with DPBS (Thermofisher), and microscopy confirmed non-astrocyte cell removal. Trypsin treatment, neutralization, and detachment were performed, and the cell suspension was transferred to tubes. Additional washes, centrifugation, and resuspension in serum-free DMEM were carried out. Cell counting utilized a trypan blue solution on a Countess slide. Astrocytes were then re-plated onto three 1.5H poly-D-lysine coated glass bottom petri dishes and three glass NWs substrates and cultured for 7 days, with½ media change every 3 days. Cells were fixed on day 7, which involved 15 minutes of incubation at room temperature with 4% paraformaldehyde (PFA, Biotium) followed by three washes with DPBS for 5 minutes each.

### 2.5. Astrocyte purity assessment

The characterization of astrocytes involved immunostaining for glial fibrillary acidic protein (GFAP), a commonly used marker of astrocytes. The astrocytes were fixed with 4% PFA for 20 minutes at room temperature, followed by three washes with DPBS for 5 minutes each. Fixed cells were then permeabilized with 0.2% Triton X-100 (Sigma-Aldrich) for 30 minutes at room temperature, and then blocked with blocking buffer composed of DPBS with 5% normal goat serum (Thermofisher), 0.2% Triton X-100, and 1% bovine serum albumin (BSA, Sigma-Aldrich) for 1 hour at room temperature. Then, cells were incubated with GFAP (1:1000, Dako), diluted in blocking buffer overnight at 4°C. After the incubation, cells were washed with DPBS for 10 minutes three times and incubated for an hour with the secondary antibody, Alexa Fluor 488 (1:1000, Thermofisher), diluted in the blocking buffer at 4°C. Astrocytes were also stained with Hoechst 33342 (1:2000, Thermofisher) and washed three times with DPBS for 5 minutes each and kept in dark until imaging.

### 2.6. Optical diffraction tomography (ODT)

ODT was performed to capture astrocyte morphology on the glass NWs and glass substrates using the HT-X1 system from Tomocube. This system is comprised of a 40x, 0.95 NA air objective, and a 450nm LED light source. 3D RI tomogram images and 2D fluorescence images were captured. TomoStudio (TomoCube, Republic of Korea) was used to visualize and obtain 3D RI tomogram images and 2D maximum intensity projection (MIP) images.

### 2.7. Morphological analysis of ODT images

Astrocyte primary process is defined as the process that extends directly from the soma. Other processes that split from the primary processes are considered branchlets. The process lengths and soma area were calculated using the TomoAnalysis software (TomoCube, Republic of Korea). The total arbor length is defined as the sum of all the primary processes in a cell. The distance of the intersection to the nucleus was calculated by first fitting the nucleus to an ellipse and finding the center, and then measuring the distance of each intersection of processes to the center of the ellipse. Cell segmentation was performed manually on 2D MIP images using ImageJ. After cell segmentation, we obtained between around 80 astrocytes per sample type. These 2D segmented cells served as masks and were applied to the stacks of 3D RI tomogram images to generate 3D RI tomograms featuring only one cell per field of view [40]. Typically, the original 3D RI tomogram images contain multiple cells. Quantitative analyses were performed on 3D RI tomogram images with one cell per field of view using MATLAB. Area was calculated by counting the number of nonzero pixels in the masks. Cell dry mass values were obtained by initially setting refractive index thresholds and then calculating the surface integral of the optical path difference over the specific refractive index increment [41]. Volume was calculated by counting the number of nonzero pixels in 3D segmented images.

## 3. Results and Discussion

### 3.1. From silicon nanowires to glass nanowires: Fabrication, morphology, optical and mechanical properties

Transparent nanostructured substrates composed of SiO_2_ nanowires (referred to as glass NWs) were synthesized via thermal oxidation of silicon nanowires (Si NWs). Initially, Si NWs were grown through plasma enhanced chemical vapor deposition (PECVD) on fused silica substrates, resulting in a sample with an opaque and brown colored appearance, as depicted in the photograph of **Figure 2a**. Afterwards, the Si NWs underwent thermal treatment in an oxygen atmosphere at temperatures reaching 980°C overnight: the oxygen diffused into the Si NWs and reacted with Si forming SiO_2_ in amorphous phase [35], [36], [37]. Thus, we obtained the transparent and weakly reddish sample of the photograph in **Figure 2b**. Scanning electron microscopy (SEM) images of these structures, as grown and after thermal treatment, are presented in **Figures 2c** and **2b**, respectively. The images revealed a dense ensemble of disordered and randomly oriented NWs with a tapered shape in both cases. Comparative analysis of the length and diameter distributions in **Figures 2f** and **2e** showed comparable lengths spreading in the range 1-3 μm. However, glass NWs exhibited significantly larger diameters, ranging from 80-180 nm, compared to Si NWs’ diameters of about 50-80 nm. These morphological modifications originated from the large difference in molar volumes between Si and SiO_2_, resulting in volume expansion of the nanowires when the Si NWs were transformed into glass NWs [42]. Interestingly, these volume modifications did not cause breaking or damaging of the NWs, as evidenced by comparing the SEM images obtained from Si NWs and glass NWs.

**Figure 2.**
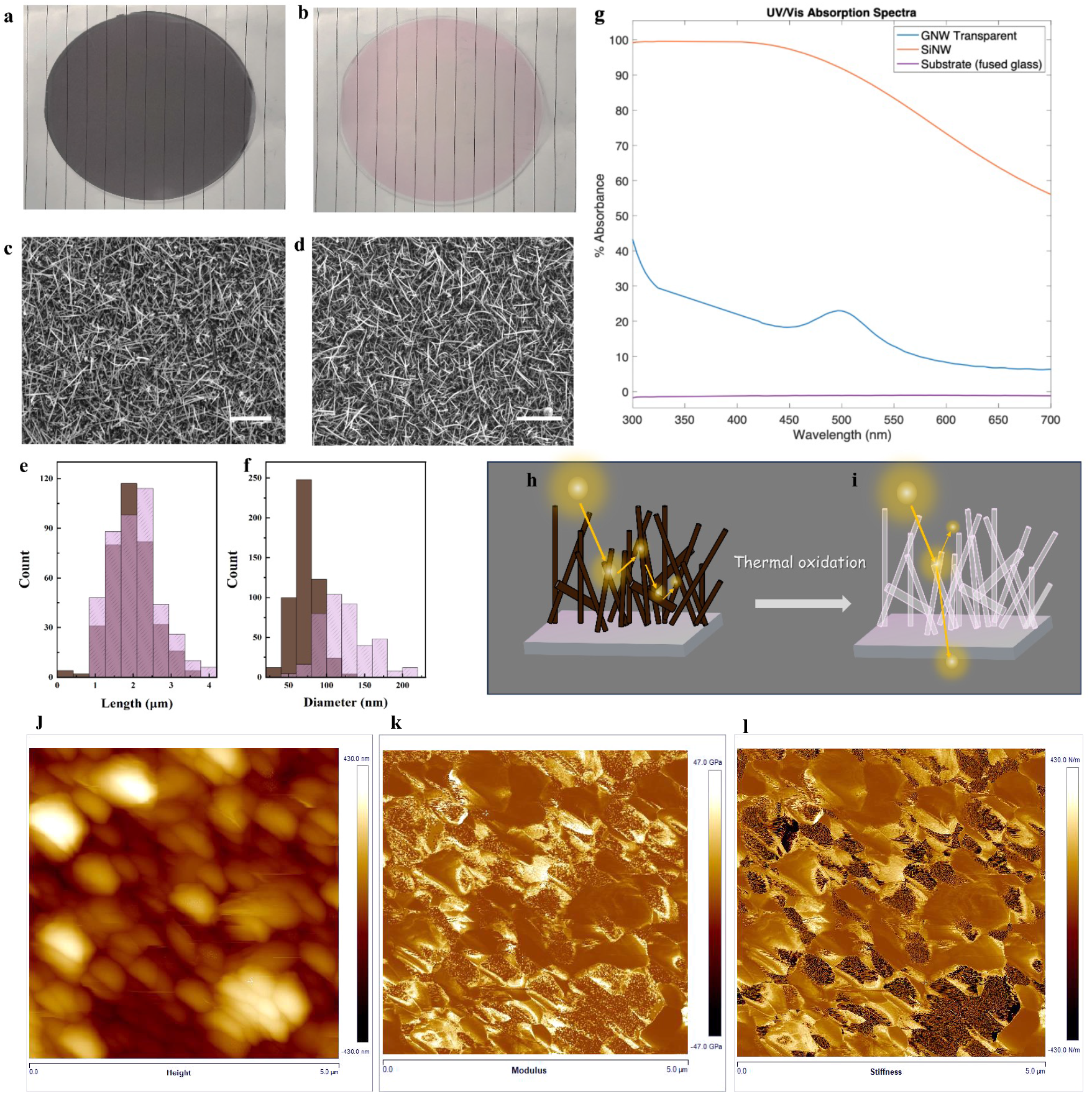
Photographs of the Si NWs on silica wafer before (a) and after thermal oxidation, referred to as glass NWs (b). SEM images of the as-grown Si NWs (c) and glass NWs (d), scale bars are 5 μm. Length (e) and diameter (f) distributions for the as-grown Si NWs (brown histograms) and glass NWs (pink histograms). Absorbance spectra, A(λ), of as-grown Si NWs (brown line) and glass NWs (magenta line) compared with bare substrate (black line) (g). Illustrations of light propagation in the Si NWs mat (h), showing the multiple scattering mechanism, and in the glass NWs mat (i), behaving like a continuous and transparent thin film. (j-l) AFM data showing the modulus and stiffness ranges on one region of the glass NWs substrate.

To quantitatively characterize the optical behavior, we conducted transmittance (T) and reflectance (R) measurements across the spectral range of 300-700 nm. **Figure 2g** compares the absorbance spectra, defined as A(λ)=100-T(λ)-R(λ), of the as-grown Si NWs (brown line), the glass NWs (magenta line), and the bare fused silica substrate (black line). The A(λ) spectra clearly indicate: i. the Si NWs mat was a strongly absorbing material; ii. glass NWs mat was characterized by a remarkable decreased absorbance, as low as 20% over the entire analyzed range, with the occurrence of an absorption peak at the wavelength of about λ=500 nm, imparting the reddish appearance observed in **Figure 2b**.

The optical behavior of Si NWs mat can be considered as results of the light trapping, a phenomenon common in semiconductor nanowires [43], [44]: the light passing through a Si NWs mat, like sketched in Figure 1h, undergoes multiple scattering events from each NW when the condition *d*_*NW∼*_*λ/10* occurs, being d_*NW*_ the later size of NWs and λ the incident light wavelength. In our case, Si NWs size and tapered shape assured indeed a large light scattering for all UV-visible wavelengths. Thus, the scattering events fold the light path many times in a random walk inside the mat causing an enhancement of the light absorption in the absorption spectral range above the electronic band gap of the Si (i.e., for wavelengths λ<1.1 μm), leading to the observed high absorbance and dark-brown appearance.

The behavior observed in glass NWs can be explained by considering the optical properties of SiO_2_ that exhibits high transparency in the UV-visible spectral range, extending down to wavelengths of approximately 200 nm, with a refractive index typically ranging from 1.54 to 1.45 in the transparent region of spectrum [45]. The weak difference in refractive index between the glass NWs and the surrounding medium, such as air, results in a significantly reduced amount of light being reflected by the individual nanowires after a scattering event. Consequently, light propagates through the mat of glass NWs as if through a continuous and transparent medium, like sketched in Figure **2i**. In addition we should consider that the growth of Si NWs is based on the vapor liquid solid mechanism that needs the presence of the Au nanoparticles acting as catalysts to favor the nucleation of Si NWs [34], [46], [47]. In our case, we deposited on fused silica wafer a 2 nm thick Au layer that dewetted at the growth temperature of 350°C by generating nanoparticles. Thus, the absorption peak observed in the absorbance spectrum of glass NWs (**Figure 2g**) is ascribable to the plasmonic resonance of the Au catalysts. Eventually, to complete the material characterization we performed AFM measurements to evaluate the glass NWs mechanical characteristics (**Figures 2j-l**). It was found a modulus range of about 94 GPa and a stiffness range of about 860 N/m.

### 3.2. Quantitative Morphological Analysis of Advanced maturation Astrocytes on Glass Nanowires

Astrocytes were cultured on glass NWs substrates and on plain glass (control) for 7 days to study the influence of substrate topography on astrocyte morphology. We acquired ODT images from astrocytes cultured on both glass NWs substrates (**Figure 3a**) and glass (**Figure 3b**) substrates. Remarkable differences in astrocyte morphology were observed between those cultured on glass (**Figure 3b**) versus glass NWs (**Figure 3a**) substrates. Astrocytes cultured on traditional glass substrates typically displayed a flattened and rounded morphology, while those cultured on glass NWs substrates exhibited a striking “star-like” shape, characterized by multiple processes extending from the cell body. A deeper analysis of the morphology of astrocytes cultured on glass NWs allows us to distinguish the primary processes, which are the main branches extending directly from the cell soma, and branchlets, such as secondary and tertiary processes, extending from these primary branches (**Figure 3a, Figure 4a**) [48], [49]. A cell with only two primary processes is intermediate maturation, whereas a cell with more complex branching indicates an advanced maturation state.

**Figure 3.**
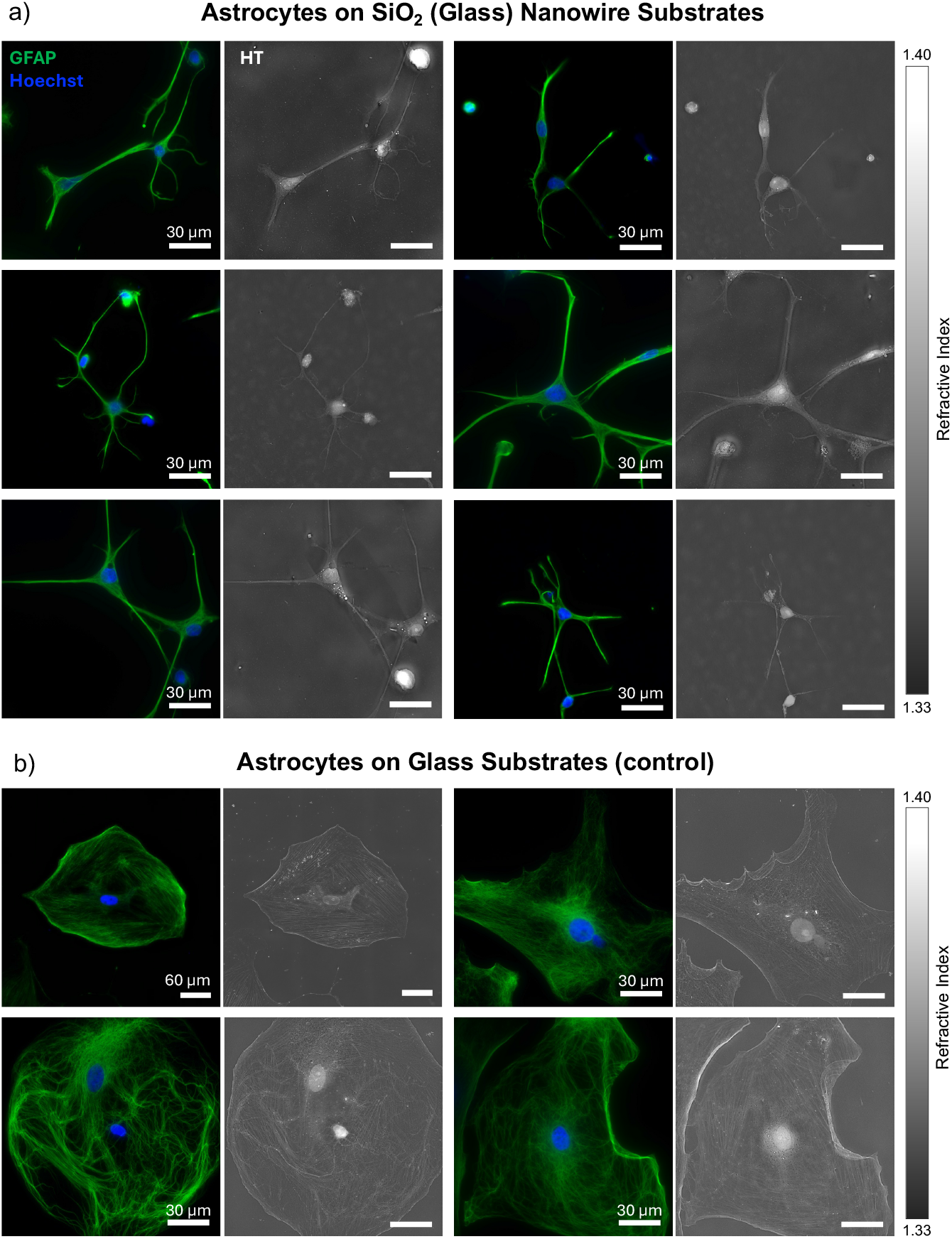
ODT images of day 7 astrocytes on SiO_2_ NWs substrates (a) and on glass (b). Fluorescence images show glial fibrillary acidic protein (GFAP) (green) and nucleus (blue) expressions along with the corresponding maximum intensity profile (MIP) images to the right of them. Bars represent the range of RI values.

**Figure 4.**
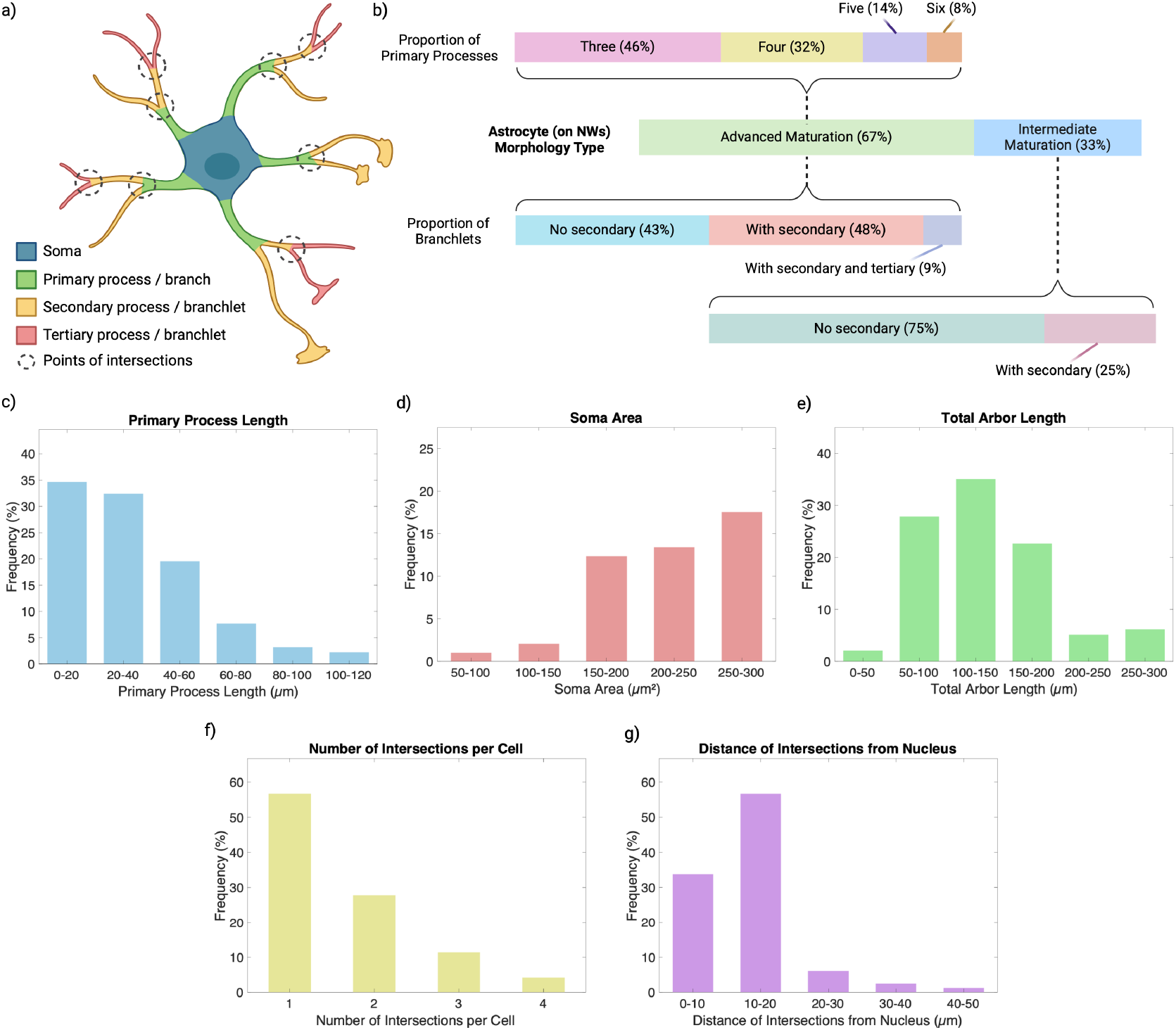
Analysis of processes of astrocytes cultured on glass NWs substrates. (a) schematic indicating parts of the advanced maturation astrocyte. (b) proportion of intermediate maturation (2 processes) vs advanced maturation (3-6 processes) astrocytes, proportion of primary processes for advanced maturation astrocytes, and proportion of branchlets. (c) distribution of primary process lengths, (d) soma area, (e) primary process total arbor length, (f) intersections per astrocyte, and (g) distance from intersection to nucleus for astrocytes cultured on glass NWs substrates.

**Figure 4** presents a detailed analysis of advanced maturation astrocyte processes cultured on glass NWs substrates. The main objective of these analyses is to demonstrate that culturing astrocytes on glass NWs substrates promotes the development of complex, advanced maturation astrocyte morphology. In this study, astrocytes with up to six processes were observed, which shows that glass NWs substrates support the development of these complex morphologies. Additionally, ODT enabled visualization of secondary and even tertiary levels of branching. Remarkably, over half of the astrocytes cultured on glass NWs had at least secondary branchlets, emphasizing the ability of this substrate to promote advanced morphological advanced maturation. ODT images also allowed for the analysis of intersections per cell and the distance from each intersection to the nucleus, which is a standard metric in astrocyte morphology studies for several reasons. Mainly, it helps quantify the complexity of the astrocyte’s branching structure, which is important for understanding how astrocytes interact with their environment.

**Figure 4b** highlights the proportion of intermediate maturation versus advanced maturation astrocytes. Approximately 33% of the astrocytes are classified as intermediate maturation, having only two processes extending from the soma, while the majority, 67%, are advanced maturation, with 3 to 6 processes. Among the advanced maturation astrocytes, the majority (46%) possess three processes, followed by 32% with four processes, 14% with five, and 8% with six processes. The presence of multiple processes indicates increased branching complexity. Additionally, it shows that 43% of astrocytes lack secondary branchlets, while 48% possess secondary branchlets, and a small percentage (9%) even demonstrate tertiary branchlets, further emphasizing their complex morphology. The ability of glass NWs to support astrocytes with up to six processes implies that the substrate design plays a significant role in the outcome of astrocyte morphology, suggesting that NWs provide a structural support not achievable on flat glass (control) surfaces. This advanced maturation-promoting quality of NWs could be harnessed to create more physiologically relevant models for CNS research.

**Figure 4 (c–g)** reveals key morphological trends in astrocytes cultured on glass NW substrates. The primary process length (**Figure 4c**) predominantly falls within the 0–40 µm range, with the highest frequency in the 0–20 µm range. Data aligns with known principles of cellular response to micro- and nano-topographies. Studies have shown that topographical cues, like those provided by NWs, play a significant role in cytoskeletal organization which directly impact process extension and cellular morphology [50]. Astrocytes interacting with a structured topography often display longer processes due to enhanced mechanotransduction, which triggers signaling pathways that promote cytoskeletal rearrangement and cellular extension [51]. *In vivo* intermediate maturation processes are linked with mature astrocyte phenotypes that facilitate connections with multiple synapses and blood vessels, enhancing astrocytic functionality [52]. Thus, the observation that advanced maturation astrocytes exhibit longer primary processes on NW substrates promote these NWs as inducers of maturation and advanced maturation. The soma area (**Figure 4d**) shows a more even distribution, with the highest frequency in the 200–250 µm^2^ range. Smaller soma areas (<150 µm^2^) and larger ones (>250 µm^2^) are less frequent, demonstrating that somas of most astrocytes fall within a moderate size range.

For the total arbor length (**Figure 4e**), the majority of astrocytes cultured on glass NWs substrates have lengths in the 100–200 µm range, peaking at 100–150 µm. This is a direct reflection of the enhanced branching capacity of advanced maturation cells, facilitating the formation of complex networks that are critical for astrocyte functions in synaptic support and homeostasis. **Figure 4f** highlights that around 90% of astrocytes exhibit at least three intersections. Both the arbor length and intersections that lead to further branching are important, as astrocytes expand their branching networks to form intricate connections within the brain’s cellular environment. For example, this is especially critical for maintaining synaptic homeostasis, facilitating metabolic exchange in the CNS and forming the tripartite synapse structure, where astrocytes contribute to neurotransmitter uptake and ionic regulation [53], [54].

The distance of intersections from the nucleus (**Figure 4g**) reveals that most intersections occur within 20 µm of the soma. The increased distance of intersections from the nucleus further underscores the biological importance of these morphological changes. In mature astrocytes, distal branching allows for more extensive coverage of synapses, enhancing their ability to regulate extracellular ion concentrations, remove neurotransmitters, and participate in tripartite synapses, where astrocytes, neurons, and synaptic structures interact. The ability to form such networks is vital for proper brain function and plasticity, especially in response to neuronal signaling [30], [55], [56].

The morphological distinctions observed in **Figure 4 (c–g)** astrocytes cultured on glass NWs substrates provide important insights into the cellular changes associated with astrocyte maturation and their role in network formation. As highlighted in the introduction, astrocytes play a crucial role in supporting neuronal function by forming intricate networks that maintain homeostasis and facilitate synaptic activity [30], [32], [57]. The results presented here demonstrate that advanced maturation of astrocytes, induced by nanoscale cues from glass NWs substrates, is accompanied by significant changes in cellular morphology that reflect their evolving biological functions. Overall, **Figure 4** underscores the ability of glass NWs substrates to promote advanced maturation of astrocytes and the development of complex branching structures. The star-like morphology and the tertiary level of branching observed in a significant portion of the cultured astrocytes indicate that the substrate effectively mimics an *in vivo*-like environment, making it an ideal platform for studying astrocyte behavior and function. The hypothesis supported by the data is that NW substrates may also facilitate astrocyte mechanotransduction by inducing cellular signaling cascades involved in cytoskeletal rearrangement and process extension.

In **Figure 5a**, the mass of astrocytes cultured on glass NWs substrates is found to be comparable to that of the control group, indicating consistent growth. **Figures 5b** and **5c** reveal exciting insights, showing that astrocytes on glass NWs demonstrate higher density while exhibiting slightly lower volume compared to those on traditional glass. This suggests a more efficient cellular composition and organization fostered by the interaction with the substrate. Additionally, **Figure 5d** highlights area measurements, where astrocytes on glass NWs occupy a larger area than their counterparts in the control group, further supported by the ODT images in **Figure 3**. This illustrates the profound impact of substrate structure on cellular morphology. Overall, the data presented in **Figure 5** strongly indicate that the unique properties of glass NWs positively influence astrocyte composition and organization through substrate interactions. This emphasizes the vital role of substrate design in enhancing astrocyte behavior and paves the way for future studies.

**Figure 5.**
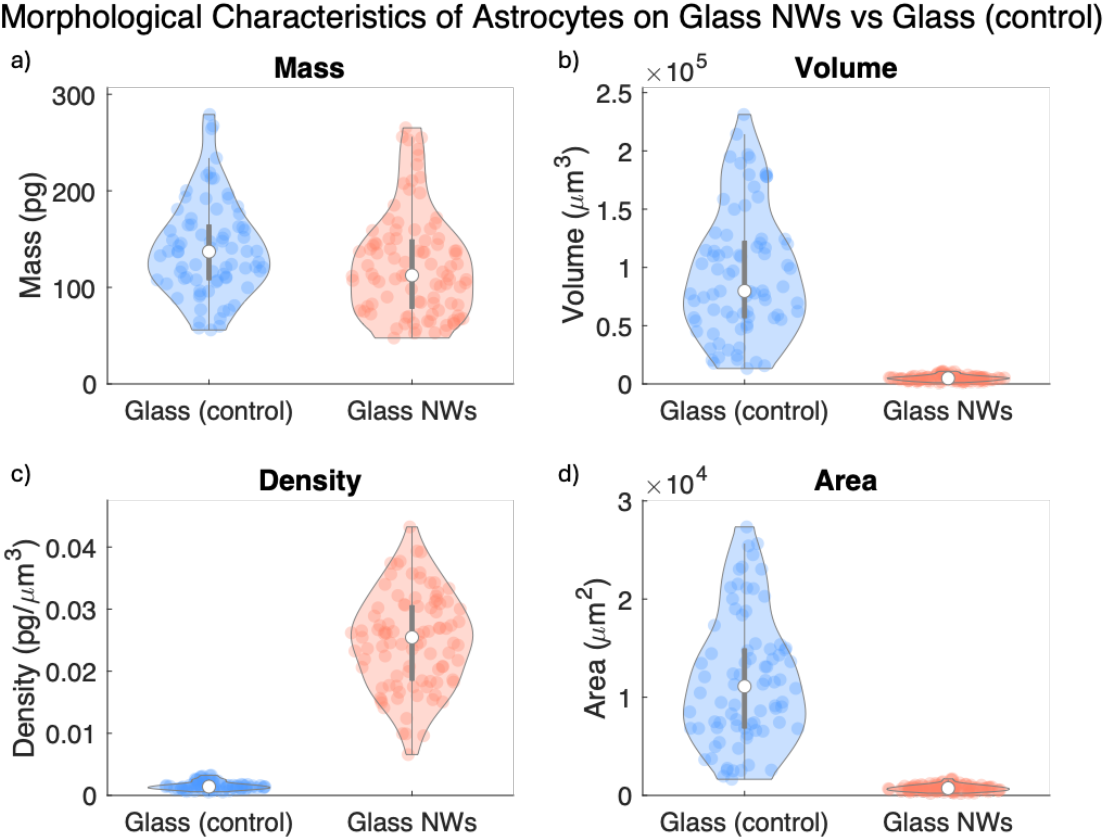
Morphological characteristics of astrocytes cultured on glass NWs vs glass (control). Box plots show comparison for (a) mass, (b) volume, (c) density, and (d) area.

In summary, the utilization of glass NWs substrates provides a biomimetic environment that mimics the physiological conditions experienced by astrocytes *in vivo*. This encourages the cells to adopt a morphology that closely resembles their natural shape, facilitating more accurate observations and analyses of their behavior and interactions with the substrate. Furthermore, ODT represents a breakthrough in imaging technology by enabling quantitative assessment of cellular structures with high precision. It allows for the quantitative measurement of optical phase delay maps and outputs 3D RI distribution of the sample. By harnessing the capabilities of ODT, we can delve deeper into the intricate cell-material interactions, shedding light on fundamental aspects of neurobiology such as cell migration, communication, and response to external stimuli.

## 4. Conclusion

Since we have established that astrocytes cultured on glass NWs exhibit a star-like morphology typically found in vivo, this substrate presents a promising model for future studies. The glass NWs mimic the textured ECM, providing a more physiologically relevant environment compared to traditional flat glass or plastic substrates. This is significant because it allows astrocytes to behave more naturally, as demonstrated by their complex branching patterns and advanced maturation. The ability of these nanostructured surfaces to promote such *in vivo*-like morphology suggests that glass NWs are highly suitable for culturing astrocytes, making them an ideal platform for further research on cell-ECM interactions.

A major significance of this study is that it serves as a pilot, laying the foundation for more in-depth investigations by integrating glass NWs substrates with ODT. ODT, an advanced imaging technique, enables the quantification of cellular characteristics with picogram precision, providing detailed insights into subtle changes in cell morphology, growth, and advanced maturation. By capturing these dynamics in 3D, ODT reveals structural information that would be challenging to obtain with conventional 2D imaging techniques. We have demonstrated that astrocytes thrive on this substrate, as evidenced by their *in vivo*-like, star-like morphology and the intricate tertiary branching that indicates an advanced maturation state. This advanced maturation has been readily captured and analyzed using ODT. Our findings highlight the potential of glass NWs substrates as nanostructured supports for ODT imaging, providing a new approach for future studies aimed at understanding astrocyte function in the CNS, as well as their roles in neurodevelopment and neurodegenerative diseases.

Moreover, glass NWs substrates could be employed in studies requiring the preservation of native cell morphology for accurate results, such as drug screening or therapeutic research. This platform provides a unique opportunity to study how the physical environment influences astrocyte behavior and to further explore astrocyte-neuron interactions in a more realistic setting. As this is only the beginning, future research could investigate how varying the architecture of the nanowires or combining them with other materials might influence cellular processes. The results from this study demonstrate that glass NWs are an excellent model for culturing astrocytes and suggest numerous applications in neurobiology.

## Acknowledgments

Support for this work was provided by CNR-JHU Joint Laboratory “Integrating transparent nanowires in optical diffraction tomography to investigate collective cell behaviour”, and from the Air Force Office of Scientific Research (FA9550-22-1-0334).

Figure 1 and Figure 4a-b was created using www.biorender.com.

